# Two Ubiquitin-Activating Systems Occur in Plants with One Playing a Major Role in Plant Immunity

**DOI:** 10.1101/2021.09.02.458739

**Authors:** Bangjun Zhou, Chaofeng Wang, Xuanyang Chen, Yi Zhang, Lirong Zeng

**Affiliations:** Plant Science Innovation Center and Plant Pathology Department, University of Nebraska, Lincoln, NE 68588; Biology Department, University of Arkansas, Little Rock, AR 72204

## Abstract

Many plants possess two or more ubiquitin-activating enzymes (E1). However, it is unclear whether the E1s of a plant genome play equivalent roles in various pathways. Here we report that tomato and tobacco encode dual ubiquitin-activating systems (DUAS) in which the E1s UBA1 and UBA2 display differential specificities in charging four groups of E2s. The C-terminal ubiquitin-folding domain of the E1s play a major but not sole role in determining the differential specificities of charging the four groups E2s. The dual systems do not play equivalent roles in plant immunity, with silence of UBA2 only compromising host immunity. Among the differentially charged E2s, group IV members UBC32, UBC33 and UBC34 are shown to be essential for ER-associated protein degradation (ERAD) and plant immunity. Like tomato, Arabidopsis UBC32/33/34 E2 triplet are also differentially charged by its E1s and are essential for plant immunity. Loss of function in Arabidopsis UBC32, UBC33 and UBC34 does not affect flg22 and elf18-triggered inhibition of seedling growth but results in alteration of ER stress tolerance, which likely contribute to the diminished plant immunity in the mutants. Our results uncover DUAS in plants and a previously unknown E1–ERAD-associated E2 triplet module in the regulation of host immunity.

**One sentence summary:** Plant dual ubiquitin E1 systems play distinct roles in plant immunity by differentially charging the ERAD-associated E2s for ER stress tolerance.

## Introduction

Ubiquitination is a post-translational protein modification (PTM) that plays key roles in numerous cellular and physiological processes. The stepwise enzymatic cascade catalyzing ubiquitination typically consists of three different classes of enzymes, ubiquitin-activating (E1), ubiquitin-conjugating (E2), and ubiquitin ligase (E3) (Callis, 2014). In the E1-E2-E3 cascade, E1s stand at the apex thus modification of proteins by ubiquitin depends on the abundance, activity, and specificity of the E1 enzymes (Schulman and Harper, 2009).

Humans possess dual E1 activation systems for ubiquitin that are directed by two distantly-related E1 enzymes UBE1 and UBA6 (Jin et al., 2007). The UBA6 and UBE1 display distinct preferences for E2 charging *in vitro*, with the E1-E2 specificity depending partly on their C-terminal Ufd domain, which is similar to that of the yeast E1 (Jin et al., 2007; Lee and Schindelin, 2008; Olsen and Lima, 2013). The UBA6 orthologues were detected in vertebrates and the echinoderm sea urchin but not in insects, worms, fungi, and plants (Jin et al., 2007). In plants, homologs of human UBE1 have been isolated with ubiquitin-activating activity being demonstrated from wheat (Hatfield and Vierstra, 1992), *Nicotiana tabacum* (Takizawa et al., 2005), Arabidopsis (Hatfield et al., 1997) and soybean (Zhang et al., 2018). While most plant species encode two or more E1s, it remains unknown whether the plant E1s have different specificities for E2 charging. Neither has been elucidated whether and how different ubiquitin E1s from a given plant genome play different roles. In *N. tabacum*, expression of the two E1 genes, *NtUBA1* and *NtUBA2* was induced in response to viral infection, wounding, and defense-related hormones, leading to the speculation that they might play equal roles in stress responses (Takizawa et al., 2005). However, the two Arabidopsis E1 enzymes apparently function differentially in plant responses to biotic stress (Goritschnig et al., 2007).

In eukaryotes, endoplasmic reticulum-associated protein degradation (ERAD) is part of the ER-mediated protein quality control (ERQC) machinery, which includes chaperone-mediated assistance in protein folding and the selective degradation of terminally misfolded proteins by ERAD (Buchberger et al., 2010; Li et al., 2020). The terminally misfolded proteins are first recruited by adapters such as the HRD3 protein, to different ER membrane-anchored E3 ligase complexes, such as the HRD1 (HMGCOA Reductase Degradation 1) and the DOA10 (Degradation Of Alpha2 10) complex, followed by retro-translocation, ubiquitination, and subsequent 26S proteasome-dependent degradation in the cytoplasm (Buchberger et al., 2010; Strasser, 2018). Failure to remove misfolded proteins by ERAD is associated with more than sixty human diseases, including neurodegenerative diseases, diabetes, and cancer (Guerriero and Brodsky, 2012). ERAD has also emerged in recent years to play an important role in modulating plant responses to biotic and abiotic stress (Chen et al., 2020). In Arabidopsis, the E2 enzyme AtUBC32 serves as an active ERAD component and functions in the salt stress tolerance (Cui et al., 2012b). Although the E2 enzymes AtUBC33 and AtUBC34 were also shown to be localized to the ER membrane (Ahn et al., 2018), evidence for the involvement of AtUBC33 and AtUBC34 in ERAD is lacking. To date, no ERAD-associated E2 enzymes have been reported to be involved in plant immunity.

In this study, we found that both tomato (*Solanum lycopersicum*) and *Nicotiana benthamiana* possess <u>d</u>ual E1 <u>u</u>biquitin-<u>a</u>ctivating <u>s</u>ystems (DUAS) that are directed by two E1s, UBA1 and UBA2. The dual systems involve differential charging of four groups E2s and do not play equivalent roles in plant immunity and development. The C-terminal ubiquitin-folding domain to the tomato E1s were shown to play a major but not sole role in governing differential charging of the four groups E2s. Among the E2s that are differentially charged by the E1s are the group IV that consists of homologs to the Arabidopsis E2s AtUBC32, AtUBC33 and AtUBC34. The tomato and *N. benthamiana* UBC32, UBC33 and UBC34 are shown to be essential for ERAD and plant innate immunity. Noteworthy, AtUBC32, AtUBC33 and AtUBC34 are also found to be differentially charged by the Arabidopsis E1s and essential for host immunity, suggesting DUAS may be conserved in many plants. Loss of function in AtUBC32, AtUBC33 and AtUBC34 does not affect flg22 and elf18-triggered suppression of seedling growth but results in alteration of ER stress response, which likely contribute to the diminished plant immunity in the mutants. Additionally, the AtUBC32 and AtUBC33/34 appear to play differential roles in ER stress response, suggesting complexity in the modulation of plant immunity by the E1–group IV E2 triplet module.

## Results

### Tomato and Tobacco Genomes Each Encode Two Ubiquitin E1s

The E1 enzymes possess a signature architecture that contains three conserved domains, a pseudo-dimeric adenylation domain involved in the ubiquitin activation, a Cys domain harboring the catalytic cysteine residue, and a ubiquitin-fold domain (Ufd) that participates in recruitment of E2 (Lee and Schindelin, 2008; Olsen and Lima, 2013; Schäfer et al., 2014). When the sequences of Arabidopsis and wheat ubiquitin E1 enzymes were used to search the tomato genome, two genes (Solyc06g007320) and Solyc09g018450) were identified that encode proteins containing all the conserved domains of E1 and with a deduced molecular mass of ~ 110 kilodalton (Supplemental Figure S1) (Hatfield and Vierstra, 1992; Hatfield et al., 1997; The tomato genome Consortium, 2012). The two genes were named *SlUBA1* (*<u>S</u>olanum <u>l</u>ycopersicum* <u>u</u>biquitin-<u>a</u>ctivating enzyme1) (Solyc06g007320) and *SlUBA2* (Solyc09g018450), respectively in the order of being cloned. *In vitro* thioester assay (Kraft et al., 2005; Takizawa et al., 2005) showed that both SlUBA1 and SlUBA2 can catalyze formation of tomato E2 SlUBC3-ubiquitin adducts that is sensitive to DTT (Figure 1A), indicating they both possess ubiquitin-activating activity.*N. benthamiana* is a solanaceous species related to tomato and has served as an important model system for studying plant immunity. When sequences of SlUBA1 and SlUBA2 were used to search homologs in the *N. benthamiana* genome (Bombarely et al., 2012), two genes (Niben101Scf09017g00015 and Niben101Scf03202g13010) were found to encode proteins that contain all the hallmark E1 domains and to be highly homologous to SlUBA1 and SlUBA2, respectively. Consistent with earlier report (Jin et al., 2007), no homologous genes of human UBA6 were identified in the tomato and *N. benthamiana* genomes. Phylogenetic analysis indicated that SlUBA1 and SlUBA2 have the highest homology with NbUBA1 and NbUBA2, respectively (Supplemental Figure S2).

**Figure 1.**
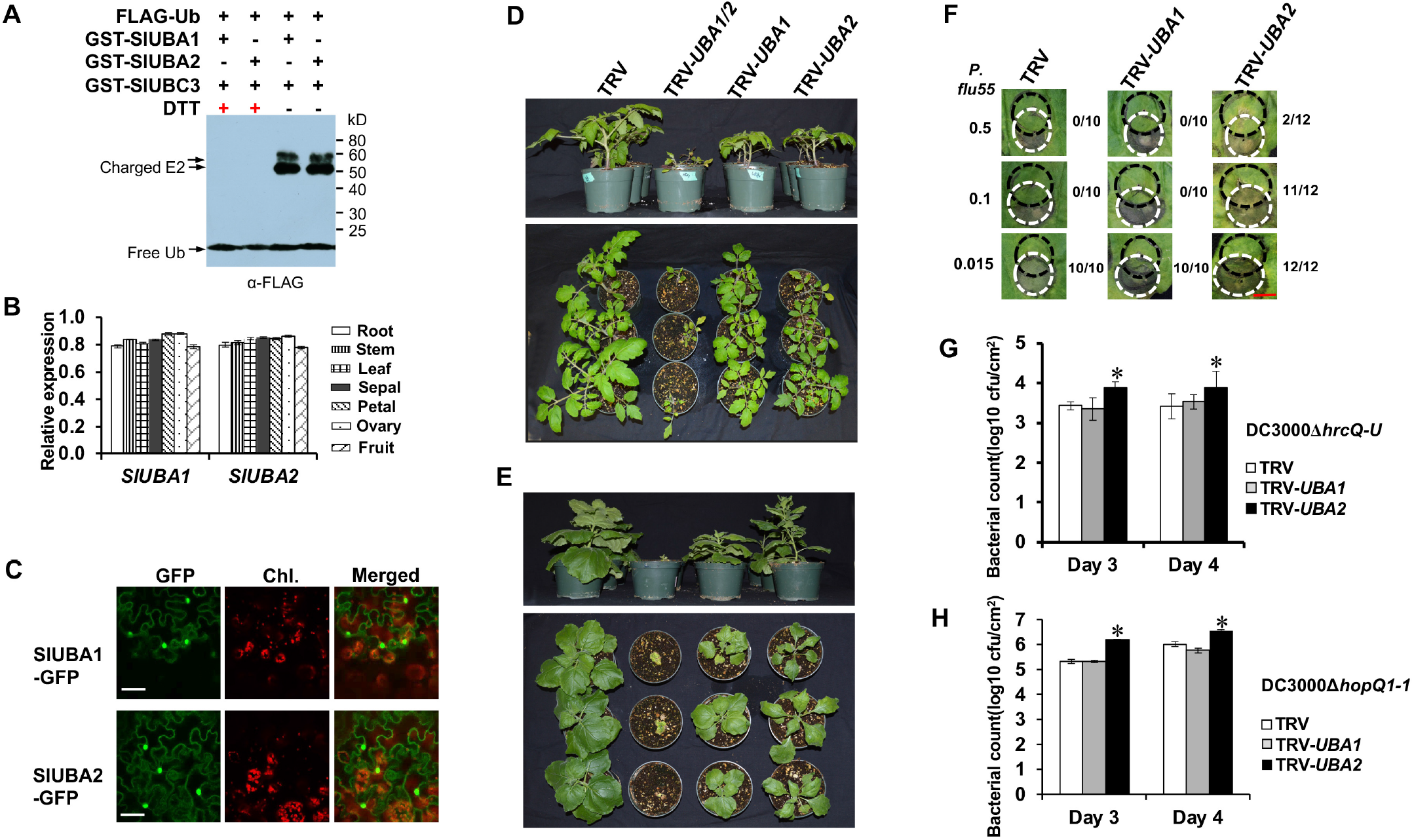
Tomato genome encodes two ubiquitin E1s that function differentially in plant development and host immunity. **(A)** The tomato proteins SlUBA1 and SlUBA2 encode active E1 enzyme. The numbers on the right denote the molecular mass of marker proteins in kD. The experiment was repeated two times with similar results. **(B)** The *SlUBA1* and *SlUBA2* genes show comparable level of expression in various tomato organs. **(C)** SlUBA1 and SlUBA2 are presented in both cytoplasm and nucleus. GFP-fused SlUBA1 and SlUBA2 in tobacco leaves was examined by confocal microscopy. Bars = 20 μm. **(D)** and **(E)** The tomato and tobacco E1s differentially regulate plant development. Tomato **(D)** and tobacco **(E)** plants in which the E1 gene *UBA1* (TRV-*UBA1*), *UBA2* (TRV-*UBA2*) or both (TRV-*UBA1/2*) are silenced are shown. The non-silenced TRV empty vector (TRV) was used as control. Up panel: side view; low panel: top view. Photographs were taken 4 weeks after the approximately 3-week-old seedlings were infiltrated with corresponding VIGS construct. **(F)** VIGS of *UBA2* gene in *N. benthamiana* compromised PTI-mediated cell death suppression. Black dashed circles denote the infiltration area of *P. fluorescens* 55 (*P. flu55*) while white dashed circles denote infilatration area of *Pst* strain DC3000. Numbers at the left side denote the corresponding concentration of *P. flu55* (OD_600_ value) used to activate PTI. Numbers at the right side of each image represent the number of overlapped infiltration areas that displayed cell death and the total number of infiltrated overlapping areas. Photographs were taken on day 4 after infiltration of *Pst* DC3000. Bar = 1 cm. **(G)** and **(H)** Bacterial growth on the *UBA1*- and *UBA2*-silenced tomato **(G)** and tobacco (*N. benthamiana*) **(H)** plants. Non-silenced (TRV) plants were used as control. Tomato VIGS Plants **(G)**were vacuum infiltrated with *Pst* strain DC3000*ΔhrcQ-U*. Tobacco VIGS Plants **(H)** were vacuum infiltrated with *P. flu55* to induce PTI and then inoculated with *Pst* strain DC3000*ΔhopQ1-1* six hours later. Experiments were repeated three times with similar results. Asterisks indicate significantly elevated bacterial growth compared to the control plants based on the one-way ANOVA (P < 0.01).

Most plant species possess two or more ubiquitin E1s (Supplemental Table I). To address whether the E1s of a given plant genome function equivalently or distinctly, we first analyzed the expression and subcellular localization of the two tomato E1 genes. Both *SlUBA1* and *SlUBA2* were expressed in all the tomato organs tested and no significant difference in the level of expression was detected between *SlUBA1* and *SlUBA2* (Figure 1B). In addition, SlUBA1 and SlUBA2 both expressed in the nucleus and cytoplasm (Figure 1C).

### SlUBA1 and SlUBA2 Play Different Roles in Plant Development and Immunity

To find out whether the two tomato E1s also function similarly, we silence the expression of the *UBA1* and *UBA2* gene in tomato and *N. benthamiana* by virus-induced gene silencing (VIGS) (Supplemental Figure S4). Based on the alignments of tomato and *N. benthamiana UBA1* and *UBA2* gene, we chose a fragment from *SlUBA1* gene for specifically silencing tomato *SlUBA1* and *N. benthamiana NbUBA1*, and a fragment from *SlUBA2* gene for specifically silencing of *SlUBA2* and *NbUBA2* (Supplemental Figure S3). These two fragments were joined together for silencing both *UBA1* and *UBA2* in tomato and *N. benthamiana*. The *UBA1* and *UBA2* genes of tomato and *N. benthamiana* were silenced specifically and efficiently in TRV-*UBA1*, TRV-*UBA2* and TRV-*UBA1/2* infected plants (Supplemental Figure S4). Silencing of *UBA1* and *UBA2* caused growth changes in tomato and *N. benthamiana* plants. Both *UBA1* and *UBA2*-silenced plants displayed reduced growth compared to the control plants (Figure 1D, 1E and Supplemental Figure S5). *UBA1*-silenced plants were dwarf with shorter nodes, a shorter taproot and fewer branch roots and slightly smaller leaves, whereas TRV-*UBA2* infected plants were nearly as tall as the control plants and a shorter taproot and significantly fewer branch roots, and smaller, narrowly shaped leaves. Tomato and *N. benthamiana* plants where both *UBA1* and *UBA2* were silenced were severely affected in growth and development, rapidly etiolated and eventually died within five to seven weeks after TRV-*UBA1/2* inoculation.

Expression of the tomato *UBA1* and *UBA2* genes is induced upon treatment of leaves with the immunogenic peptide of flagellin, flg22 (Supplemental Figure S6), suggesting they both are involved in host immunity. To find out whether the tomato and *N. benthamiana* E1s also play different roles in immunity, we employed two assays to evaluate plant pathogen-associated molecular pattern (PAMP)-triggered immunity (PTI) on *UBA1* and *UBA2-*silenced *N. benthamiana* plants. Cell death suppression assay (CDSA) was first performed on *UBA1* and *UBA2-*silenced and control *N. benthamiana* plants (Chakravarthy et al., 2010). In CDSA, PTI induced by the nonpathogen *P. fluorescens* 55 on the *N. benthamiana* plants inhibits the hypersensitive cell death induced by subsequent inoculation of *Pseudomonas syringae* pathovar *tomato (Pst)* strain DC3000 in the overlapping area (Figure 1F). However, our results showed that cell death was observed in the overlapping area on *UBA2*-silenced plants but not *UBA1*-silenced *N. benthamiana* plants, which implies a breakdown of PTI induction on *UBA2*-silenced plants. We further examined the effects of silencing *UBA1* and *UBA2* on restriction of *Pst* strains DC3000*ΔhrcQ-U* and DC3000*ΔhopQ1-1* growth on tomato and *N. benthamiana*plants, respectively (Wei et al., 2007; Zhou et al., 2017). The T3SS (type three secretion system)-deficient *Pst* strain DC3000*ΔhrcQ-U* elicits PTI on tomato plants (Kvitko et al., 2009). Growth of DC3000*ΔhrcQ-U* on the *UBA2*-silenced tomato plants was significantly higher at day 3 and day 4 after inoculation than that on the *UBA1*-silenced and TRV-infected control plants (Figure 1G). Pre-inoculation with the nonpathogen *P. fluorescens* 55 would induce PTI and enhance differences in pathogenic bacterial growth between wild-type and PTI-defective *N. benthamiana* plants (Nguyen et al., 2010). Accordingly, the growth of *Pst* DC3000Δ*hopQ1-1* on the *UBA2*-silenced plants was significantly higher than that on the *UBA1*-silenced and non-silenced control plants at day 3 and day 4 after inoculation (Figure 1H). Thus, our results confirmed that tomato and *N. benthamiana* UBA1 and UBA2 function differentially in host immunity.

### Tomato Possesses DUAS in which UBA1 and UBA2 Differentially Charge Four Groups of E2s

E1s initiate the enzymatic cascade for ubiquitination by coordinating ubiquitin activation with transferring to cognate E2s. To find out the underlying molecular basis for differential roles of tomato E1s in host immunity, we examined the specificities of tomato UBA1 and UBA2 in charging of a panel of 34 tomato E2s (Zhou et al., 2017). The majority of the E2s tested can be charged by both SlUBA1 and SlUBA2 at comparable specificities (Figure 2A and Supplemental Figure S7, Supplemental Table II). By contrast, E2s in groups IV, V, VI, and XII were charged by SlUBA2 at significantly higher efficiency. In particular, the group V E2s were efficiently charged by SlUBA2 but not charged (SlUBC7 and SlUBC36) or at extremely lower efficiency (SlUBC14 and SlUBC35) by SlUBA1. Thus, tomato possesses dual E1 activation systems for ubiquitin. Interestingly, the tomato group IV, V, and VI E2s that are differentially charged by SlUBA1 and SlUBA2 are phylogenetically close to those human E2s that are also differentially charged by human E1s UBE1 and UBA6 (Supplemental Figure S8).

**Figure 2.**
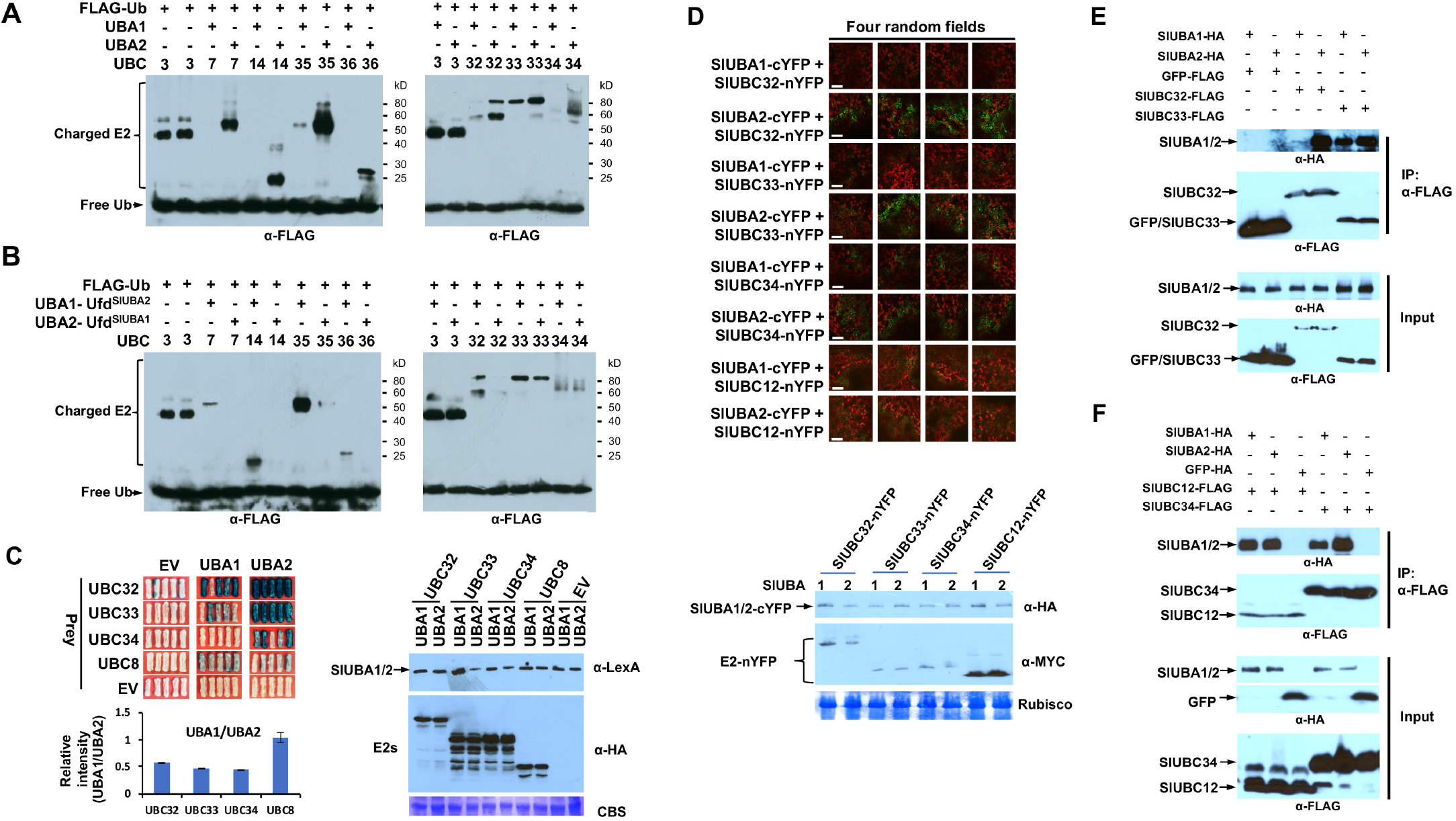
The tomato E1s display different specificities to E2s *in vitro* and *in vivo*. **(A)** SlUBA1 and SlUBA2 differentially charge groups IV (SlUBC32, 33 and 34) and V (SlUBC7, 14, 35 and 36) E2s in thioester assay. The experiment was repeated at least three times with similar result. SlUBC3 was used as control. **(B)** SlUBA1-Ufd^SlUBA2^ and SlUBA2-Ufd^SlUBA1^ reverse the specificities of SlUBA1 and SlUBA2 in charging group IV and V E2s. The numbers on the right denote the molecular mass of marker proteins in kilodaltons. Experiments were repeated three times with similar results. (**C**) The interaction of SlUBA1 and SlUBA2 with group IV E2s was detected by yeast two-hybrid. SlUBC8 and the empty prey and bait vectors (EV) were used as control. Interaction was demonstrated by activation of the lacZ reporter gene (blue patches). Photographs were taken 24 h after streaking the yeast cells onto plates containing X-Gal. The low left panel shows ratio of blue color intensity in yeast cell patches of UBA1-E2 compared to that of the UBA2-E2 as shown in upper left panel. The right panel shows comparable level of SlUBA1 and SlUBA2 protein and similar level of the E2 protein were expressed for each pair of SlUBA1-E2 and SlUBA2-E2 in the yeast cells. CBS, Coomassie blue staining as an indicator of equal sample loading. (**D**) Tomato UBA2 showed significantly stronger interaction with SlUBC32, SlUBC33 and SlUBC34 than UBA1 in the BiFC assay. SlUBC12 was used as control. Images from four random microscopic fields for each tested construct pair were presented. The experiment was repeated two times with similar results. Scale bar = 20 μm. Low panel shows the level of protein expressed *in planta* for the E1 (UBA1 or UBA2)-E2 pairs tested in the assay. (**E**) and (**F**) Significantly higher amount of tomato UBA2 than UBA1 was pulled down by SlUBC32, SlUBC33 and SlUBC34 in co-immunoprecipitation assay. FLAG-tagged SlUBC12 and GFP were used as negative controls. The experiment was repeated two times with similar result.

To find out if the Ufd domain also plays a role in differential E2 charging by plant E1s, we exchanged the Ufd domain of SlUBA1 and SlUBA2 to build chimeric SlUBA1 and SlUBA2 proteins (SlUBA1-Ufd^SlUBA2^ and SlUBA2-Ufd^SlUBA1^) (Supplemental Figure S9A and S9B). The chimeric SlUBA1-Ufd^SlUBA2^ and SlUBA2-Ufd^SlUBA1^ proteins charged the SlUBC3 (as control) at similar efficiency, which is the same as that when SlUBA1 and SlUBA2 were used (Figure 1A). By contrast, E2s from group IV and V were charged by the chimeric SlUBA1-Ufd^SlUBA2^ at much higher efficiencies, which is opposite to the results that SlUBA1 and SlUBA2 were used. Thus, the Ufd domain of tomato E1s also plays a major role in the specificity of E2 charging. The charging of E2s by SlUBA1-Ufd^SlUBA2^ is slightly weaker than by SlUBA2 (Figure 2A), suggesting the Ufd domain may not be the sole factor that determine the E2 charging specificities. This was further supported by the result that no difference was detected in the strength of interaction between the Ufd domain of SlUBA1 (Ufd^SlUBA1^) and SlUBA2 (Ufd^SlUBA2^) and the group IV E2s (Supplemental Figure S9C).

### The SlUBA2 Has Higher Specificities Than SlUBA1 and Play A Major Role in Charging Group IV E2s *in vivo*

The distinct roles of tomato and *N. benthamiana* E1s in host immunity are likely due to their differential specificities in charging of certain members of the four groups of E2, which act with cognate E3s to target plant immune components. Considering the involvement in numerous human diseases and the emerging role in plant immunity for ERAD and the involvement of AtUBC32 in ERAD (Cui et al., 2012b; Guerriero and Brodsky, 2012), we decided to focus on the group IV E2s for further characterization of the plant DUAS. Higher specificity of an E1 in charging E2s would be manifested by stronger E1-E2 interaction, as the factors that govern affinity and specificity for a target protein are the same (Eaton et al., 1995). We thus employed three different assays of E1-E2 interaction to test the specificities of SlUBA1 and SlUBA2 for group IV E2s *in vivo*. First, the yeast two-hybrid assay showed group IV E2s have much stronger interaction with SlUBA2 (Figure 2C). And the differential interactions were not caused by differential levels of protein expression (Figure 2C, right panel). Similarly, SlUBA2 has stronger interactions with group V E2s than that of the SlUBA1 in yeast two-hybrid (Supplemental Figure S10A). Next, much stronger *in planta* interaction of SlUBC32, SlUBC33, and SlUBC34 with SlUBA2 than with SlUBA1 was detected in the bimolecular fluorescence complementation (BiFC) assay (Figure 2D). The SlUBA1 and SlUBA2 protein as well as the corresponding E2 protein in each pair of SlUBA1-E2 and SlUBA2-E2 being compared were expressed at comparable level (Figure 2D, lower panel). Lastly, co-immunoprecipitation was employed to detect the interactions of the two tomato E1s with group IV E2s. As shown in Figure 2E and 2F, group IV E2s pulled down significantly more SlUBA2 than SlUBA1 in the assay. Together, these results indicate that SlUBA2 possesses higher specificity towards group IV E2s than that of SlUBA1 and likely plays a major role in charging the E2 triplet *in vivo*. To further corroborate this conclusion, we tested charging of the group IV E2s *in planta* by transient expressing *myc*-tagged E2s on *N. benthamiana* plants where either *UBA1* or the *UBA2* genes was silenced. Compared to the control plants, no noticeable difference in charging of group IV E2 was observed on the *UBA1*-silenced plants. By contrast, silencing of *UBA2* essentially abolished the charging of the group IV E2s (Supplemental Figure S10B).

### Group IV E2s Are Required for Plant Immunity

Quantitative PCR (qPCR) indicated that group IV E2 genes, *SlUBC32, SlUBC33* and *SlUBC34* were significantly induced two hours after flg22 treatment (Figure 3A) and 24 hours after inoculation of the tomato plants with *Pst* strains DC3000 (Figure 3B). The *SlUBC33* and *SlUBC34* were also induced at 24 hours after inoculation of DC3000*ΔhrcQ-U*. Growth of the *Pst* strain DC3000*ΔhrcQ-U* was significantly higher on tomato plants in which the *SlUBC32, SlUBC33* and *SlUBC34* genes were efficiently knocked down by VIGS (Figure 3C) than that on the control (TRV) plants on day 3 after inoculation (Figure 3D and Supplemental Figure S11A), suggesting that group IV E2s are required for plant immunity.

**Figure 3.**
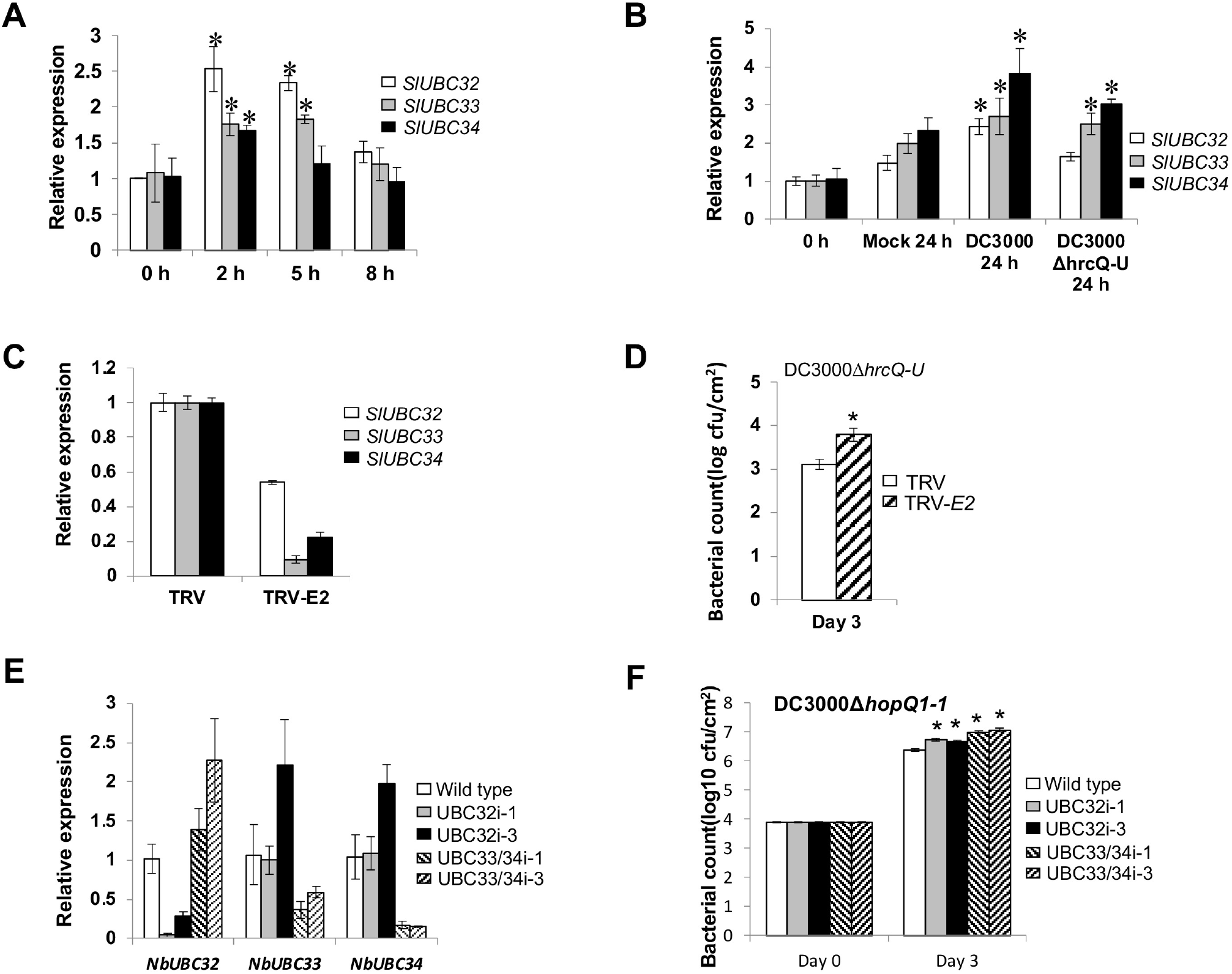
Group IV E2s are required for plant immunity. (**A**) and (**B**) The expression of tomato *SlUBC32, 33* and *34* genes was induced by flg22 (**A**) and different *Pst* strains **(B)**. The expression of *SlUBC32, SlUBC33* and *SlUBC34* were analyzed by qRT-PCR using tomato *EF1α* gene as the internal reference. The experiment was performed with three technical repeats in each of the three biological replicates. Error bars indicate standard deviation. Asterisks indicate significantly elevated expression compared to mock plants at the same time point based on the one-way ANOVA (P < 0.01). (**C**) and (**E**) The E2 genes were efficiently and specifically silenced in tomato (**C**) and tobacco (**E**) plants. The accumulation of the transcript for E2 genes was quantified by qRT-PCR. *EF1α* was used as an internal reference. In tomato (**C**), plants infected with the empty TRV vector were used as control (TRV). In tobacco (**E**), wild type plants were used as control. The relative quantity of transcript for corresponding gene in control plants was set as 1. Experiments were repeated at least two times with similar results. (**D**) Knocking down the expression of tomato group IV E2 genes significantly increased bacterial growth (DC3000*ΔhrcQ-U*) compared to the control plants infected with the TRV empty vector (TRV). The dipping method was used for pathogen inoculation. (**F**) Knocking down the expression of tobacco group IV E2 genes in transgenic plants significantly increased bacterial growth (DC3000*ΔhopQ1-1*) compared to the control plants. In (**D**) and (**F**), asterisks indicate significantly increased bacterial growth compared to control plants based on the one-way ANOVA (P < 0.01).

The *N. benthamiana* genome encodes a corresponding close homolog for each of the tomato E2s (Zhou et al., 2017). In order to confirm the roles of group IV E2s in plant immunity, we developed transgenic *N. benthamiana* lines in which group IV E2 genes are silenced. Among the group IV E2 genes, *UBC33* and *UBC34* are highly homologous but have relatively lower homology to *UBC32* (Zhou et al., 2017). Therefore, the transgenic lines UBC32i and UBC33/34i where the *UBC32* and the *UBC33*/*UBC34* gene are specifically silenced, respectively were developed (Figure 3E). Growth of the *Pst* strain DC3000Δ*hopQ1-1* on the *UBC32*-silenced (UBC32i) and *UBC33/34*-silenced (UBC33/34i) plants was significantly higher than that on the wild type *N. benthamiana* plants on day 3 after inoculation (Figure 3F). Consistently, silencing of *UBC32, UBC33* and *UBC34* in *N. benthamiana* plants by VIGS also resulted in reduced host immunity (Supplemental Figure S11B).

### Group IV E2s Are Localized to ER and Interact with ERAD-Related E3s

Similar to their Arabidopsis counterpart, tomato SlUBC32, SlUBC33 and SlUBC34 possess a conserved transmembrane domain (TMD) (Supplemental Figure S12) (Ahn et al., 2018). In Arabidopsis, ER-localized AtUBC32 interacts with the HRD1/HRD3A E3 complex to serve as part of the ERAD system (Cui et al., 2012b; Chen et al., 2016). SlUBC32, SlUBC33 and SlUBC34 fused with GFP to the C-terminus co-localized with the mCherry-fused, ER-localized Marker ER-rb (Nelson et al., 2007) in *N. benthamiana* leaves (Figure 4A), indicating they are also ER-bound. By contrast, the localization to ER was abolished when the TMD of SlUBC32, SlUBC33 and SlUBC34 was deleted. Next, we examined the interaction of group IV E2s with the tomato closest homolog of the ERAD-related E3s AtHRD1A and AtHRD1B. The tomato HRD1A (SlHRD1A) and HRD1B (SlHRD1B) are localized to the ER (Supplemental Figure S13). As shown in Figure 4B, SlUBC32, SlUBC33 and SlUBC34 all interact strongly with SlHRD1A, SlHRD1B and their adapter protein SlHRD3A in yeast two-hybrid assay using the mating-based split-ubiquitin system (mbSUS) (Grefen et al., 2009). To further confirm the association of SlUBC32, SlUBC33 and SlUBC34 with E3 SlHRD1A and SlHRD1B, the BiFC assay was performed using tomato protoplasts. SlUBC32, SlUBC33 and SlUBC34 interacted with the SlHRD1A and SlHRD1B in the assay to generate green fluorescence signals whereas no signals were detected in control (Figure 4C). These results suggest that SlUBC32, SlUBC33 and SlUBC34 are also likely involved in ERAD.

**Figure 4.**
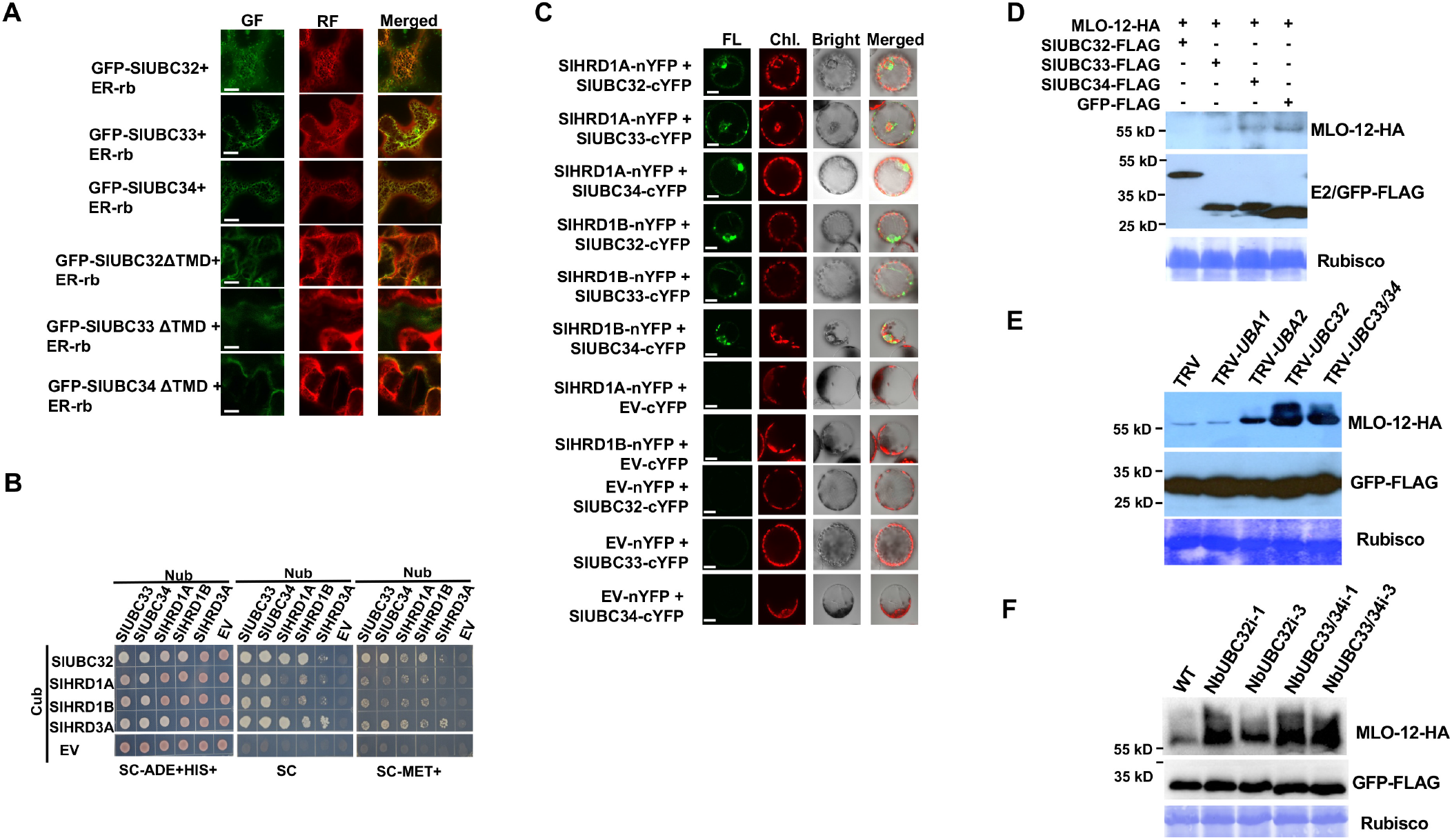
Tomato UBC32, UBC33, and UBC34 are endoplasmic reticulum (ER)-bound and are involved in ERAD. **(A)** Tomato E2s SlUBC32, SlUBC33 and SlUBC34 are localized to the ER. GFP-fused wild type and transmembrane domain-deleted (ΔTMD) mutant version of SlUBC32, SlUBC33 and SlUBC34 were examined using confocal microscope. GF, green fluorescence; RF, red fluorescence conferred by mCherry-ER-localized marker protein (ER-rb). **(B)** SlUBC32, SlUBC33 and SlUBC34 interacted with tomato homolog of ERAD-associated ubiquitin E3 ligases, SlHRD1A and SlHRD1B and their adapter protein SlHRD3A in the mating-based split ubiquitin yeast two-hybrid system. Growth of yeast cells harboring two genes that had been fused with the N-terminal (Nub) and C-terminal (Cub) halves of the ubiquitin protein, respectively on SC-ADE+HIS+, SC- and SC- with 0.15 M MET media plates, respectively, were shown in the left, middle and right panel. Empty bait and prey vector (EV) were used as control. **(C)** SlUBC32, SlUBC33 and SlUBC34 interacted with SlHRD1A and SlHRD1B in BiFC assay using tomato leaf protoplasts. EV, empty vector; FL., fluorescence; Chl., chlorophyll autofluorescence; Bright, bright field image. Scale bar = 20 μm. **(D)** Transient co-expression in *N. benthamiana* leaves of the group IV E2 and the known ERAD target MLO-12 increased degradation of MLO-12. GFP was used as control. **(E)** Knocking down UBA2 and group IV E2s in leaves of *N. benthamiana* plants by VIGS reduced the degradation of MLO-12. **(F)** Degradation of MLO-12 was reduced in *N. benthamiana* RNAi transgenic lines where group IV E2 genes was silenced. Two independent transgenic lines in which the *NbUBC32* gene and the *NbUBC33* and *NbUBC34* genes were silenced, respectively were used for the assay. The experiments in (**E**) and (**F**) were repeated two times with similar result. In (**D**), (**E**) and (**F**), the FLAG-tagged GFP was expressed together with HA-tagged MLO-12 as an internal control of efficiency in transient expression as well as control of non-misfolded protein. Staining of the Rubisco large subunit (Rubisco) by Coomassie Blue R250 denotes equal sample loading.

### UBA2 and Group IV E2s Are Required for ERAD

To confirm the involvement of group IV E2s in ERAD, we probed stability of the known ERAD substrate protein MLO-12 under conditions that group IV E2 genes were overexpressed or silenced (Muller et al., 2005; Cui et al., 2012b). Compared to the control, transient co-expression of MLO-12 and individual member of the group IV E2s promoted the degradation of MLO-12 (Figure 4D). By contrast, silencing of *UBA2*, *UBC32* and *UBC33/34* by VIGS enhanced the accumulation of MLO-12 (Figure 4E), suggesting that UBA2 and group IV E2s are required for ERAD. No increase of MLO-12 was observed in *UBA1* gene-silenced *N. benthamiana* plants, which is consistent with the results that UBA2 play a major role in charging the group IV E2s *in vivo*. Similarly, the degradation of MLO-12 was diminished in the tobacco RNAi transgenic lines UBC32i and UBC33/34i (Figure 4F). Taken together, these results confirm that group IV E2s are active components of the ERAD system and both the group IV E2s and the UBA2 are required for ERAD.

### AtUBC32, AtUBC33 and AtUBC34 Are Differentially Charged by Arabidopsis E1s and AtUBC33 and AtUBC34 Are Also Required for ERAD

To study the role of individual members of the group IV E2s in ERAD and host immunity, we attempted to knock down each member specifically. However, the extremely high homology between *SlUBC33* and *SlUBC34* makes it highly challenging to silence them individually by VIGS or RNAi-based gene-silencing. Tomato group IV E2s interact with Arabidopsis ERAD E3 AtHRD1A *in vivo* (Supplemental Figure S14). In addition, Arabidopsis E1s AtUBA1 and AtUBA2 differentially charge tomato SlUBC32 (Figure 5A). These results suggest that the function for tomato and Arabidopsis UBC32, UBC33 and UBC34 in ERAD is likely conserved. Furthermore, AtUBA1 charged AtUBC32, AtUBC33, AtUBC34, and SlUBC32 at much higher efficiencies than that of AtUBA2 (Figure 5B), indicating that Arabidopsis also possesses dual E1 activation systems for ubiquitin, with the Arabidopsis AtUBA1 being the ortholog of tomato SlUBA2. Finally, the Arabidopsis E1s AtUBA1 and AtUBA2 were also shown to be not equally required for disease resistance and a 15-bp deletion at the C-terminus of *AtUBA1* in the *mos5* mutant compromised host immunity (Goritschnig et al., 2007). The 15-bp/5-amino acid deletion in *mos5* AtUBA1 are mapped to the Ufd domain that is highly conserved among tomato, tobacco, and Arabidopsis E1s (Supplemental Figure S15). Thus, we decided to take advantage of the null mutant lines available for Arabidopsis *AtUBC32*, *AtUBC33* and *AtUBC34* gene for our study.

**Figure 5.**
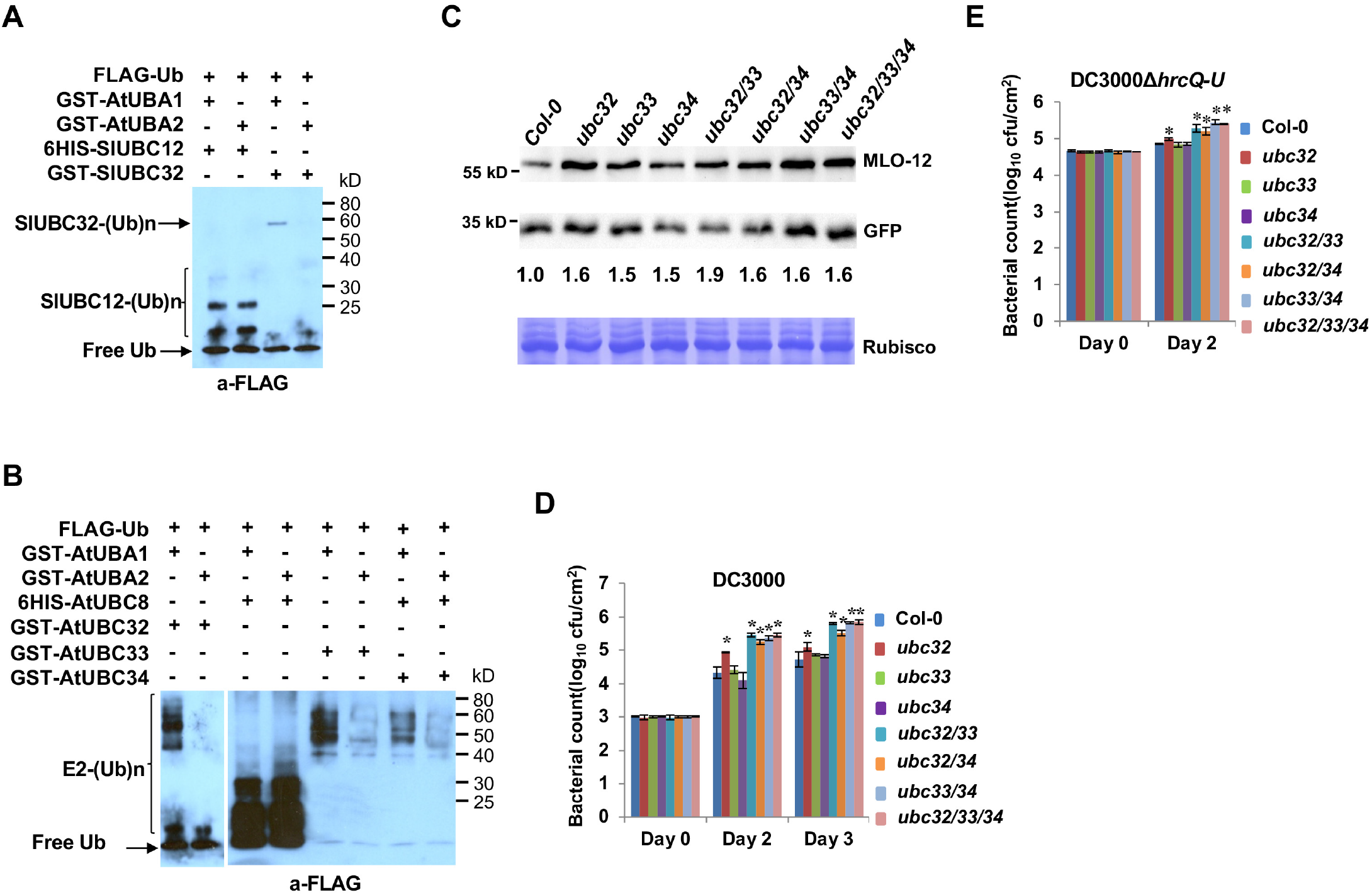
Arabidopsis E2 enzymes AtUBC32, AtUBC33 and AtUBC34 are differentially charged by the E1s AtUBA1 and AtUBA2 and are required for ERAD and host immunity. **(A)** Tomato SlUBC32 was charged by AtUBA1 with higher specificities than that of AtUBA2 in thioester assay. The tomato SlUBC12 to which tomato E1 display comparable specificities was used as control. **(B)** The Arabidopsis E2s AtUBC32, AtUBC33 and AtUBC34 were charged by AtUBA1 at significantly higher efficiencies than that of AtUBA2 in thioester assay. The AtUBC8 was used as control. The experiments in (**A**) and (**B**) were repeated two times with similar result. (**C**) Loss of function in AtUBC32, AtUBC33 and AtUBC34 diminished the degradation of MLO-12 by ERAD. MLO-12 was transiently expressed in protoplasts derived from leaves of Col-0 and the mutants, respectively. The GFP-HA was used as internal control for transfection efficiencies. **(D)**and **(E)** Bacterial growth assay on single, double and triple mutants of the Arabidopsis mutant *ubc32*, *ubc33* and *ubc34*. Wild-type Col-0 and different mutants for the *AtUBC32, AtUBC33* and *AtUBC34* gene were infected with DC3000 **(D)** and DC3000*ΔhrcQ-U* **(E)**, and then the bacterial population was examined on 2 and 3 days after infection. Bars show standard deviation. Asterisks indicate significantly elevated bacterial growth compared with the WT plants based on one-way ANOVA (P < 0.01). cfu, colony-forming units. The experiments were repeated three times with similar results.

We obtained mutant lines in which *AtUBC32, AtUBC33*, and *AtUBC34* are knocked out (Supplemental Figure S16A) and generated homozygous double and triple mutant lines *ubc32/33*, *ubc32/34*, *ubc33/34*, and *ubc32/33/34* by crossing and genotyping (Supplemental Figure S16B). The expression of the *AtUBC32, AtUBC33* and *AtUBC34* gene was examined to confirm the knock-out of *AtUBC32, AtUBC33*, and *AtUBC34* in the corresponding mutants (Supplemental Figure S16C). No significant changes in morphology are observed between Col-0 and the mutants. The *ubc33*, *ubc34*, *ubc32/33*, *ubc33/34* and *ubc32/33/34* mutant plants are slightly smaller than Col-0, whereas the mutant line of *ubc32/34* is clearly smaller than that of Col-0 (Supplemental Figure S17). The *ubc32*, *ubc33*, *ubc34* single, double, and triple mutant lines display slightly earlier flowering than Col-0 (Supplemental Figure S18).

The AtUBC32 was shown to be required for ERAD (Cui et al., 2012b). However, whether the AtUBC33 and AtUBC34 are also required for ERAD remains unknown. We expressed the ERAD substrate MLO-12 in protoplasts derived from Col-0 and Arabidopsis *ubc32*, *ubc33*, and *ubc34* mutants and monitor the accumulation of MLO-12. Consistent with previous result (Cui et al., 2012b), the null mutation in AtUBC32 diminished the degradation of MLO-12 (Figure 5C). Like AtUBC32, loss of function in AtUBC33 and AtUBC34 also reduced the turnover of MLO-12, indicating that they are involved in ERAD as well. No synergistic effect of the ubc32, ubc33 and ubc34 mutations on the degradation of MLO-12 was observed.

### Arabidopsis UBC32, UBC33 and UBC34 Are Required for Host Immunity

The mutant lines *ubc32*, *ubc32/33*, *ubc32/34*, *ubc33/34* and *ubc32*/*33/34* were more susceptible than Col-0 to *Pst* strains DC3000 and DC3000Δ*hrcQ-U*, as manifested by significantly increased pathogen growth on these lines (Figure 5D and 5E). The single mutant *ubc33* and *ubc34* displayed comparable bacterial growth to that of the Col-0 whereas the double mutant *ubc33/34* displayed significantly increased bacterial growth than Col-0, indicating functional redundancy between *AtUBC33* and *AtUBC34* in host immunity. While plants of the double mutant *ubc32/33, ubc32/34*, and *ubc33/34* are more susceptible to pathogen infection than the single mutant *ubc32*, *ubc33*, and *ubc34*, plants of the triple mutant *ubc32/33/34* displayed comparable bacteria growth as the double mutants, implying a complex relationship beyond functional redundancy and synergy of the E2 triplet in modulating plant immunity.

### Loss of Function in Arabidopsis UBC32, UBC33 and UBC34 Do Not Change flg22 and elf18-Triggered Seedling Growth Suppression but Alternate Plant ER Stress Tolerance

To elucidate the molecular underpinnings of diminished immunity in the mutant lines *ubc32*, *ubc32/33*, *ubc32/34*, *ubc33/34* and *ubc32/33/34*, we first tested whether the PRR FLS2 (Flagellin Sensing 2) and EFR (*EF*-Tu receptor)-mediated PTI are affected. FLS2 and EFR recognizes bacterial flagellin/flg22 and the elongation factor Tu/EF-Tu derived immunogenic peptide elf18, respectively to activate PTI, playing a significant role in warding off bacterial infection (Kunze et al., 2004; Zipfel et al., 2004; Yu et al., 2017). The seedling growth inhibition (SGI) assay was employed for the test, because PTI activated by flg22 and elf18 was shown to inhibit Arabidopsis seedling growth (Gómez-Gómez and Boller, 2000; Zipfel et al., 2006). As shown in Figure 6A, no significant difference in the inhibition of seedling growth was observed between the mutants and Col-0 when treated with flg22 and elf18, respectively, suggesting the FLS2 and EFR-mediated PTI was not significantly affected by loss of function in these E2 genes.

**Figure 6.**
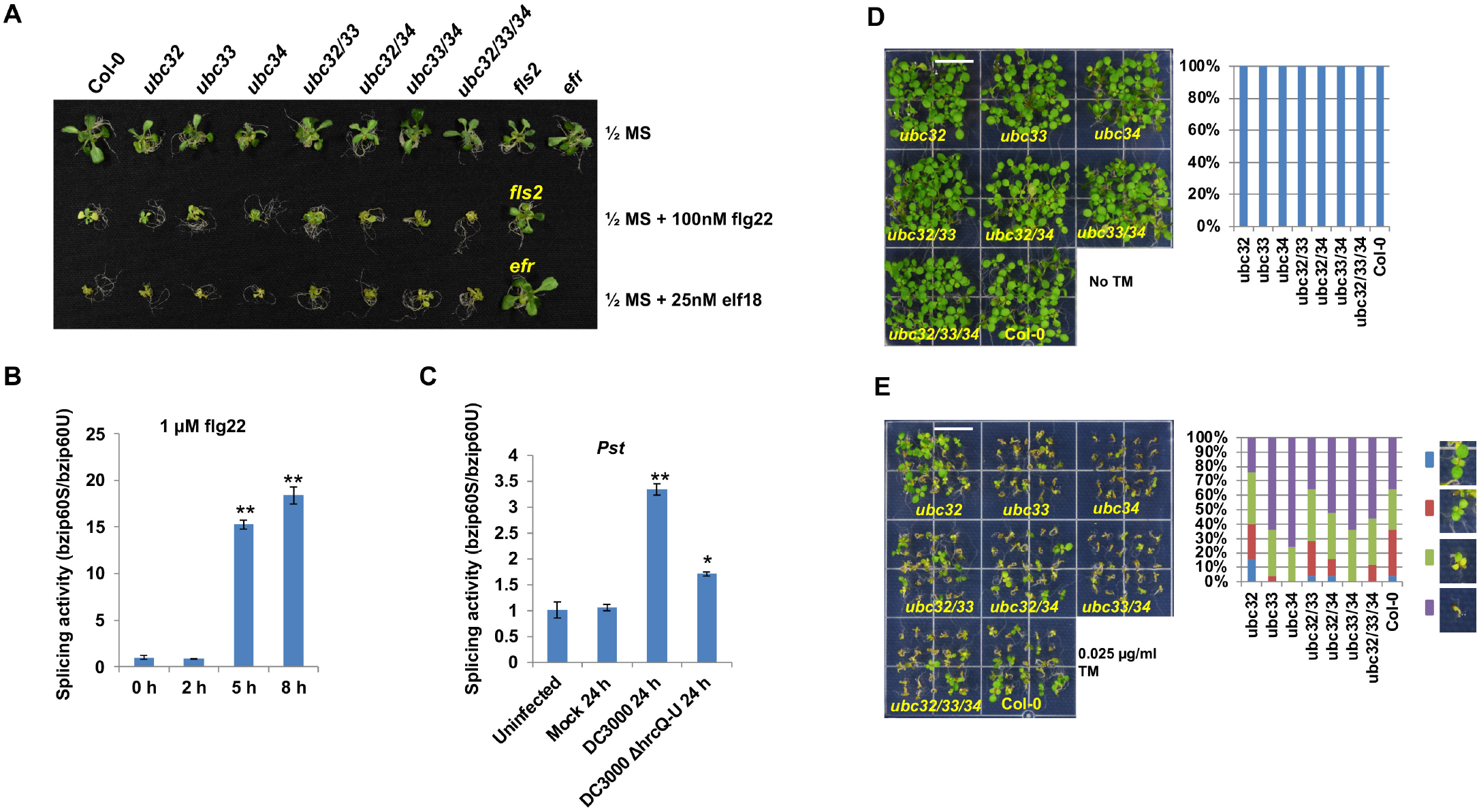
Knocking out AtUBC32, AtUBC33, and At UBC34 do not affect flg22 and elf18-triggered seedling growth inhibition but alternate plant ER stress tolerance. (**A**) Phenotype of wild-type Col-0 and *ubc32*, *ubc33*, *ubc34* single, double, and triple mutant seedlings untreated or treated with flg22 and elf18. The *fls2* and *efr* mutant were included as control. Similar results were obtained in three independent experiments. (**B**) and (**C**) ER stress response is involved in plant immunity. Quantitative measurement of bZIP60 mRNA splicing activity was performed by qRT-PCR analyses of spliced (bzip60S) and non-spliced (bzip60U) bZIP60 forms. Ratios of bzip60S to bzip60U are calculated, with setting the ratio of uninfected Col-0 as 1. (**B**) qRT-PCR analyses of bzip60S and bzip60U in 10-day tomato seedlings with 1 μM flg22 treatment in ½ liquid MS media were conducted. Ratios of spliced bzip60S and unspliced bzip60U was shown. **(C)** Ratios of spliced bzip60S and unspliced bzip60U was shown for 3-week tomato plants infected with *Pst* pathogens by dipping method. Statistical analysis was performed using Student’s t-test. *, p≤0.05. **, p≤0.001. The experiments were performed at least two times with similar results. (**D**) and (**E**) The phenotypes of different Arabidopsis *ubc32*, *ubc33* and *ubc34* mutant lines grew on media without (**D**) or with 0.025 μg/mL tunicamycin (**E**) in seedling growth assay. The percentages of different phenotypes are shown in the right panels. A representative image of two independent experiments is shown. For quantification of seedling phenotypes, n ≥ 30 (right panel). Bar = 0.5 cm. Photos were taken 7 days after sowing cold stratification-processed seeds on the growth media.

Consistent with the SGI assay, the FLS2 accumulation in the mutant lines was not significantly changed compared to that in Col-0, though *ubc32/33* has slightly decreased whereas *ubc34*, *ubc32/34, ubc33/34* and *ubc32/33/34* display slightly increased accumulation of FLS2 (Supplemental Figure S19A).

We then test whether ER stress tolerance are altered in the mutants. ER stress response has been shown to be involved in plant immunity (Moreno et al., 2012; Chakraborty et al., 2020). Indeed, both flg22 treatment and pathogen infection induce ER stress response on tomato plants, as is manifested by IRE1 (inositol-requiring enzyme 1)-mediated splicing of the ER stress response gene bZIP60 (Figure 6B and 6C). Consistent with the notion that damage to ERAD would lead to increased misfolded and unfolded proteins, the expression of the UPR marker gene *Bip3* in the mutants was enhanced compared to that in Col-0 upon treatment with the ER stress agent, tunicamycin (TM) (Supplemental Figure S19B). Seedling growth assay indicated that the *ubc32* mutant displayed higher ER-stress tolerance than that of Col-0 (Figure 6D and 6E), which is consistent with previous report (Cui et al., 2012b). The *ubc33* and *ubc34* mutant both displayed significantly reduced tolerance to TM-induced ER stress, though the *ubc33*/*34* double mutant did not show synergistic effect in reduction of ER stress tolerance. The *ubc32/33, ubc32/34* and *ubc32/33/34* displayed reduced ER-stress tolerance than Col-0, suggesting that knocking out of *UBC33* and/or *UBC34* in the *ubc32* background suppressed the elevated ER-stress tolerance caused by the *ubc32* mutation. Similarly, transgenic *N. benthamiana* 32RNAi lines displayed elevated ER-stress tolerance, whereas 33/34RNAi lines displayed reduced ER-stress tolerance (Supplemental Figure S19C). These results suggest that loss of function in UBC32, UBC33 and UBC34 results in changes of plant ER stress tolerance, which likely contribute to their role in plant immunity.

## Discussion

E1 enzymes govern the state of ubiquitination in an organism by coordinating the activation of ubiquitin with recruitment and charging of E2s, which in turn controls the downstream cognate E3s and substrates involved. The vital importance of E1s (and ubiquitination) for plants is demonstrated by the result that the tomato and tobacco plants died a few weeks after both E1 genes, *UBA1* and *UBA2* were silenced (Figure 1D, 1E and Supplemental Figure S5). Despite the importance, studies of plant ubiquitin E1s have been very limited, with essentially no functional characterizations. In the current research, we found that both tomato and *N. benthamiana* encode two ubiquitin E1s and the two E1s do not play equal roles in plant immunity and development, which can be attributed to differential charging of a subset of E2s by the E1 enzymes. While the two tomato E1s displayed comparable specificities to many E2s, including the groups III and IX E2s that have been previously shown to be required for plant immunity (Mural et al., 2013; Zhou et al., 2017; Zhou and Zeng, 2017), UBA2 charged E2s of group IV, V, VI, and XII with significantly higher specificities. When UBA1 is knocked down/mutated, the UBA2 can take up the duty of UBA1 to charge the E2s to which UBA1 and UBA2 possess comparable specificities. Therefore, charging of E2s in the cell is not affected and plant immunity is not compromised when UBA1 is silenced. However, when UBA2 is knocked down/mutated, charging of E2s in the group IV, V, VI, and XII is significantly reduced/abolished, which in turn will affect their cooperation with cognate E3s to ubiquitinate corresponding plant immunity-related substrates. Consequently, plant immunity is compromised when UBA2 is silenced (Figure 7). We also demonstrated that members of the group IV E2s, UBC32, UBC33, and UBC34 are required for ERAD, ER stress tolerance, and modulating of host immunity. Thus, our findings establish the connection between the E1-E2 module, ERAD, and host immunity.

**Figure 7.**
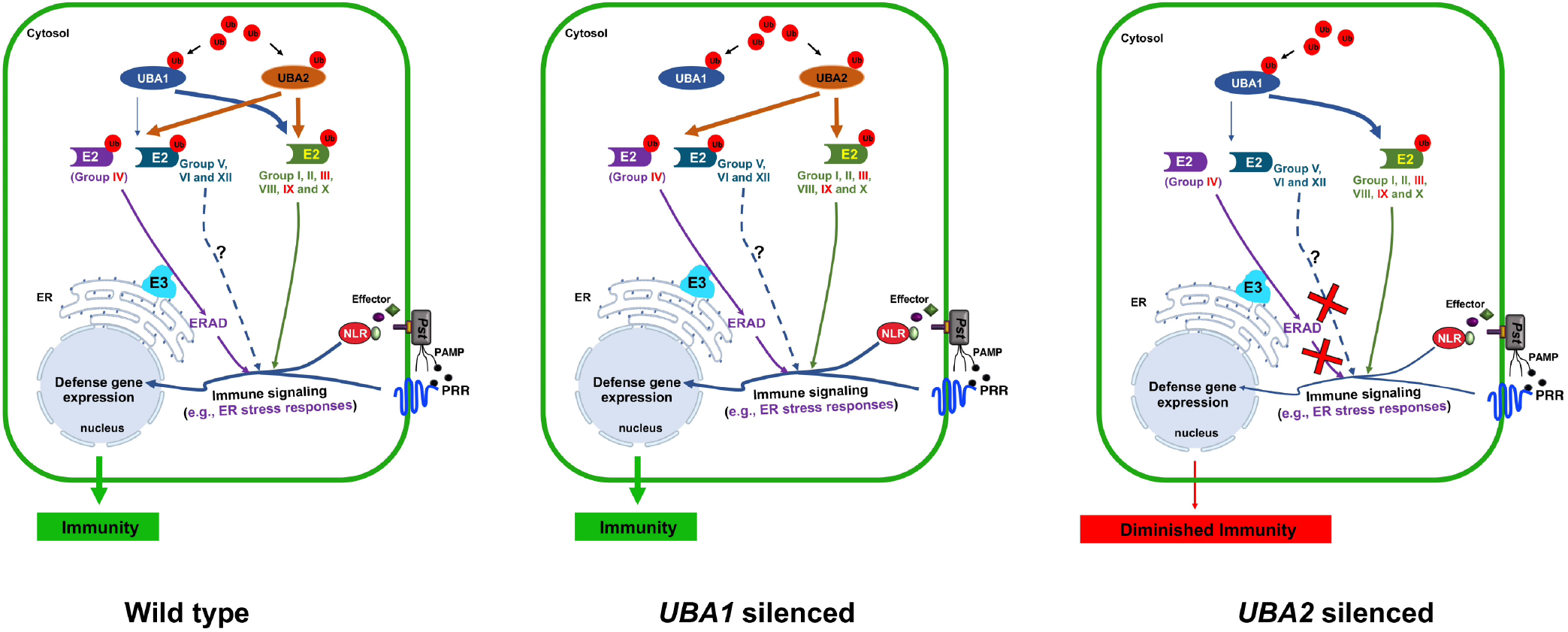
A working model illustrates how the plant DUAS function distinctly in host immunity. The two tomato E1s display comparable specificities to E2s of groups I, II, III, VIII, IX and X. However, UBA2 charges E2s of groups IV, V, VI, and XII with significantly higher specificities, which contribute to the distinct roles of UBA1 and UBA2 in plant growth, development, and defense. In the absence of UBA1, UBA2 can charge the E2s of group I, II, III, VIII, IX and X efficiently and the charging of groups IV, V, VI, and XII E2s is also not affected. By contrast, knocking out UBA2 will significantly reduce the charging of group IV, V, VI, and XII E2s, which in turn will affect cooperation of them with cognate E3s, such as working with ERAD-related E3s by group IV E2s, to modulate cognate plant immunity-related substrates by ubiquitin. Consequently, plant immunity is diminished. Nearly all sequenced plant genomes encode two or more ubiquitin E1s. Thus, plants likely encode dual or multiple E1 activation systems for ubiquitin that do not play equivalent roles in various physiological processes. Fonts in red color denotes that group III, IV, and IX E2s have been shown to modulate plant immunity. The question mark indicates that the role of group V, VI and XII E2s in plant immunity is unclear. Ub, ubiquitin; NLR, nucleotide-binding leucine-rich repeat protein; *Pst*, *Pseudomonas syringae* pv. *tomato*.

Besides group IV, SlUBA2 also has much higher specificities than SlUBA1 for charging E2s in groups V, VI and XII. Conceivably, the distinct roles of the plant DUAS in plant immunity and other physiological processes, such as plant development may be attributed to differential charging of E2s in groups V, VI and XII as well. In particular, phylogenetic analysis revealed that the tomato group V E2s fall into the same subfamily as human E2 UBE2G2 (Zhou et al., 2017). The UBE2G2 and its yeast homolog, UBC7p have been demonstrated to be involved in ERAD (Chen et al., 2006; Berner et al., 2018), which raises the possibility that the function of group V E2s are also ERAD-associated and the UBA2 of tomato and *N. benthamiana* and AtUBA1 of Arabidopsis play a major role in charging ER-related E2s. Characterization of the E1-E2 modules involving groups V, VI, and XII E2s will address the question and uncover their roles in different pathways.

Despite being closely related phylogenetically, tomato SlUBA1 and SlUBA2 differentially charge 12 out of the 34 E2s been tested. In addition, Arabidopsis E1s AtUBA1 and AtUBA2 also differentially charge the ERAD-associated E2s AtUBC32, AtUBC33 and AtUBC34, in which Arabidopsis AtUBA1 function as the ortholog of tomato SlUBA2. Thus, Arabidopsis also possesses DUAS and differential E2 charging by the dual systems appears to be conserved in plants. It has been shown that wheat encode three ubiquitin E1s, TaUBE11, TaUBE12, and TaUBE13, but TaUBE11 and TaUBE12 are 99.2% identical in amino acid sequences (Hatfield and Vierstra, 1992), leading to the speculation that wheat also encode dual E1 activation systems for ubiquitin. Considering that nearly all sequenced crop and model plant genomes encode two or more ubiquitin E1s (Supplemental Table I), plants in general likely encode dual or multiple E1 activation systems for ubiquitin that do not play equivalent roles in various physiological processes.

In addition to intrinsic specificities in charging various E2s by the ubiquitin E1, other factors, such as PTMs that affect the activities and localizations of E1s and E2s might also contribute to the distinct roles of the plant dual E1 activation systems for ubiquitin. In mammalian cells, S-glutathionylation was reported to suppress E1 and E2 activity (Jahngen-Hodge et al., 1997). Human E1 enzyme UBE1 and E2 enzymes were found to be phosphorylated *in vitro* and *in vivo* (Kong and Chock, 1992; Cook and Chock, 1995; Stephen et al., 1996). The human Casein kinase 2 (CK2) phosphorylates E2 Cell Division Cycle 34 (CDC34) to regulate its subcellular localization (Block et al., 2001). In addition, the import and/or retention of human ubiquitin E1 UBE1 to the nucleus is cell cycle-dependent (Stephen et al., 1996). More recently, structural study of the yeast E1-E2 (UBC15) complex suggests that phosphorylation of residues at the N termini of ubiquitin E2s broadly inhibits their ability to function with ubiquitin E1 (Lv et al., 2017). Although no plant E1 and E2 enzymes for ubiquitination have been shown to be modified by other PTMs thus far, it is believed that such modifications do exist in plants (Zhang and Zeng, 2020), which may serve as an extra layer of modulation in addition to differential E2 charging that lead to distinct roles of the plant DUAS.

Our findings indicate that the UBC32, UBC33 and UBC34 E2 triplet function in ERAD and plant immunity, which is conserved in tomato, tobacco and Arabidopsis. Thus, in addition to abiotic stress (Cui et al., 2012b; Ahn et al., 2018), UBC32, UBC33 and UBC34 also function in biotic stresses. The null mutants for UBC32, UBC33, and UBC34 do not display significant morphological changes, which suggest that alterations of host immunity in the mutants are unlikely caused by changes in growth and development and their role in abiotic and biotic stress pathways are specific. Noteworthy, although the enhanced bacterial growth in UBC32 knocking-out Arabidopsis mutant *ubc32* and tobacco knocking-down UBC32i plants was significant compared to the control plants, the enhanced level of pathogen growth in *ubc32* and UBC32i plants is always lower than that of the Arabidopsis *ubc33/ubc34* mutant plants and the tobacco UBC33/34i plants, respectively (Figure 3F, 5D and 5E), suggesting that UBC33 and UBC34 together contribute more to plant immunity than UBC32. The diminished plant immunity in the *ubc33/ubc34* mutant plants is in line with the result that the *ubc33* and *ubc34* mutant display reduced ER stress tolerance, as ER stress tolerance have been shown to be required for plant immunity (Wang et al., 2005; Moreno et al., 2012). By contrast, the *ubc32* mutant displays diminished immunity yet elevated ER stress tolerance, which suggests ER stress tolerance is not the sole factor that contribute to the role of UBC32 in host immunity. Indeed, previous reports indicated that UBC32 as an ERAD component negatively modulate ER stress tolerance, salt tolerance yet positively regulate oxidative burst tolerance (Cui et al., 2012a; Cui et al., 2012b). It is conceivable that the UBC32, UBC33 and UBC34 E2s can function in ERAD to target substrates of different pathways and the combination of UBC32, UBC33, and UBC34 actions at ER determine the outcome of the E2 triplet mediated ERAD in modulating plant immunity. Our results demonstrated that loss of function in the E2 triplet do not significantly affect FLS2 and EFR-mediated immunity, suggesting FLS2 and EFR may not be a key target of the E2-triplet-mediated ERAD in the modulation of host immunity. Previous studies showed that loss of function in the ER protein folding-related genes significantly affect biogenesis of EFR, whereas have merely marginal effect on that of FLS2 (Li et al., 2009; Nekrasov et al., 2009; Saijo et al., 2009). Thus, the ER apparently facilitates the folding but not degradation in the biogenesis of EFR. There are usually dozens of E2s and hundreds of E3s in a given plant genome (Vierstra, 2003; Zhou et al., 2017). Conceivably, loss of function in UBC32, UBC33 and UBC34 E2 triplet is believed to affect their cooperation with multiple cognate E3s and consequently, the modification of many substrates that reside in various pathways. It is possible that the E1-UBC32/33/34 E2 triplet module work with different E3s to modify different substrates that reside in abiotic and biotic stress pathway, respectively. In this regard, uncover and characterization of the cognate E3s and key substrates of the UBC32/33/34 E2 triplet mediated ERAD and ER stress signaling in the context of plant immunity would be the key to an in-depth understanding of the regulation of host immunity by ERAD.

## Materials and Methods

### Growth of Bacteria and Plant Materials

*Agrobacterium tumefaciens* strains GV3101 and GV2260 and strains of *Pseudomonas syringae* pv *tomato (Pst)* and *Pseudomonas fluorescens* 55 were grown at 28°C on Luria-Bertani and King’s B medium, respectively, with appropriate antibiotics. *N. benthamiana* and tomato RG-pto11 (*pto11/pto11, Prf/Prf*) seeds were germinated and plants were grown on autoclaved soil in a growth chamber with 16 h light (~ 300 μmol/m^2^/s at the leaf surface of the plants), 24°C/23°C day/night temperature, and 50% relative humidity. Arabidopsis mutant lines SALK_082711 (*ubc32*) (Cui et al., 2012b), SALK_104882C (*ubc33*) and CS878883 (*ubc34*) (Ahn et al., 2018) were obtained from ABRC. All Arabidopsis plants were grown in a growth chamber with 16 h light (~ 300 μmol/m^2^/s at the leaf surface of the plants), 22°C/22°C day/night temperature, and 50% relative humidity.

### DNA Manipulations and Plasmid Constructions

All DNA manipulations were performed using standard techniques (Sambrook and Russell, 2001). A detailed methodology is described in Supplemental Methods S1.

### Sequence Alignment and Phylogenetic Analysis

For sequence alignment, sequences of interest in the FASTA format were entered into the ClustalX 2.1 program and aligned using the ClustalX algorithm (Larkin et al., 2007). The phylogenetic analysis was then performed with the MEGAX program using the aligned sequences (Tamura et al., 2013). To build an unrooted phylogenetic tree using MEGAX, the evolutionary history was inferred using the neighbor-joining method with 1000 bootstrap trials. The evolutionary distances were computed using the p-distance method in which the evolutionary distance unit represents the number of amino acid (or nucleotide) substitutions per site (Nei and Kumar, 2000). Branches corresponding to partitions reproduced <50% bootstrap replicates were collapsed in the tree.

### Expression and Purification of Recombinant Proteins

GST-tagged fusion proteins were expressed in *E. coli* strain BL21 (DE3) and purified with Glutathione Sepharose 4 Fast Flow beads (GE Healthcare) by following the protocol provided by the manufacturer. The purified proteins were further desalted and concentrated in the protein storage buffer (50 mM Tris-HCl pH8.0, 50 mM KCl, 0.1 mM EDTA, 1 mM DTT, 0.5 mM PMSF) using the Amicon Centrifugal Filter (Millipore). The desalted and concentrated recombinant protein was stored at −80 °C in the presence of a final concentration of 40% glycerol until being used. The concentration of purified protein was determined using protein assay agent (Bio-Rad).

### Examination of Charging Ubiquitin E2s by E1s via Thioester Assay

To examine the efficiencies of charging E2s by E1s, the thioester assay was performed as described with modifications (Mural et al., 2013). In a 15 μL reaction, 40 ng of ubiquitin E1 (tomato E1 GST-SlUBA1, GST-SlUBA2, tomato chimeric E1 GST-SlUBA1-Ufd^SlUBA2^, GST-SlUBA2-Ufd^SlUBA1^, Arabidopsis GST-AtUBA1, and GST-ATUBA2, respectively) was pre-incubated with 2.0 μg of FLAG-ubiquitin in 20 mM Tris-HCl pH 7.5, 10 mM MgCl_2_, and 1 mM ATP at 28 °C for 10 min, which is followed by adding 100 ng of the GST or 6HIS - fused E2 protein to be tested. The reaction was then continued for 15 min before being stopped with SDS sample loading buffer (62.5 mM Tris-HCl pH 6.8, 2% SDS, 0.01% bromophenol blue, 10% glycerol and 4M Urea). To test the DTT sensitivity of E2-ubiquitin linkage in the thioester assay, the reaction volume was scaled up to 20 μL. The reaction was then equally split and terminated by addition of SDS sample loading buffer with either 100 mM dithiothreitol (DTT+) or 4 M urea sample buffer (DTT-). The reactions were immunoblotted with mouse monoclonal anti-FLAG M2-peroxidase-conjugated antibody (Sigma-Aldrich) before being detected using ECL kit (Pierce, now Thermo Fisher). The formation of DTT-sensitive ubiquitin adducts to tomato E2 SlUBC3 is denoted as charged E2.

### Quantitative real time-PCR (qRT-PCR)

For detecting gene expression, samples of tomato root, stem, leaf, sepal, petal, ovary and green fruit from 10-week-old tomato plants; Leaf tissues of 3 to 4-week old tomato RG-pto11 (*pto11/pto11, Prf/Prf*) plants infiltrated with 2 μM flg22 or sterilized H_2_O (mock, used as control); leaf tissues of tobacco E2-RNAi transgenic lines and VIGS plants; and leaf tissues from Arabidopsis plants with different treatments were collected for total RNA extraction using the RNeasy Plant Mini Kit with DNase treatment (QIAGEN) by following the protocol provided by the manufacturer. The first-strand cDNA was synthesized using the Superscript III reverse transcriptase and oligo dT primer (Life Technologies) according to the instructions from the manufacturer. Quantitative real-time PCR (qRT-PCR) was performed using gene specific primers and SYBR Green (Life Technologies) on the LightCycler® 480 Instrument II (Roche). All primers used in qRT-PCR are showed in the Supplemental Table III. *SlEF1a, NbEF1a* and *AtActin2* were used as the internal references for tomato, *N. benthamiana* and Arabidopsis samples, respectively.

### Yeast Two-Hybrid

For testing the interaction of two proteins using the LexA-based yeast two-hybrid system, procedures were followed as described (Golemis et al., 2008). For detecting interaction of HRD1A, HRD1B and HRD3A with UBC32, UBC33 and UBC34 using the mating-based split-ubiquitin system (mbSUS), the experiment was performed as described (Grefen et al., 2009). A detailed methodology is described in Supplemental Methods S1.

### Bimolecular fluorescence complementation (BiFC) Assay

The BiFC assay that is based on split yellow fluorescent protein (YFP) was used to test the interaction of various E1-E1, E1-E2, and E2-E3 pairs in the leaves and protoplasts by following the protocol as described (Zhou et al., 2017). A detailed methodology is described in Supplemental Methods S1.

### Coimmunoprecipitation

The coimmunoprecipitation assay of HA-tagged E1s and FLAG-tagged E2s was performed as described previously (Zhou et al., 2017). A detailed methodology is described in Supplemental Methods S1.

### Seedling growth assay

To test FLS2 and EFR-mediated immunity in Arabidopsis *ubc32*, *ubc33*, and *ubc34* single, double and triple mutant lines, seedling growth inhibition assay was performed as described (Wierzba and Tax, 2016). Specifically, Arabidopsis seeds were sterilized and then sowed on ½ MS (Murashige and Skoog) agar plates for stratification for 3 days, followed by moving the plates to short day conditions (22 °C, 10 h light, and 65-70% relative humidity) for germination for 3 days. The seedlings of Col-0 and different mutants were then transferred to liquid or solid (by adding agar) ½ MS media with or without 100 nM flg22 or 25 nM elf18 for 7 days before photographing. To detect the ER-stress tolerance of different Arabidopsis mutant lines and tobacco RNAi transgenic lines, sterilized Arabidopsis and tobacco seeds were scattered in the ½ MS plate with tunicamycin (Tm) at 0.025 (for Arabidopsis seeds) or 0.03 (for tobacco seeds) μg/mL. The development of Arabidopsis and tobacco seedlings was observed and photographed one or two weeks after sowing. Four kinds of leaf phenotypes (Green, light green, light yellow and yellow) for Arabidopsis seedlings and three kinds of leaf phenotypes (Green, light yellow and yellow) for tobacco seedlings were statistically calculated for the relative ER-stress tolerance.

### Virus-Induced Gene Silencing (VIGS)

Gene silencing was induced using the tobacco rattle virus (TRV) vectors (Caplan and Dinesh-Kumar, 2006) as previously described (Mural et al., 2013). *Agrobacterium* (OD600 = 0.5) containing appropriate pTRV plasmids was induced with acetosyringone and used to infiltrate two leaf-stage tomato seedlings and 3-week-old *N. benthamiana* seedlings. VIGS-treated tomato plants were maintained for 3 to 4 weeks at 21°C /21°C, 16/8 h day/night condition, whereas VIGS-treated *N. benthamiana* plants were maintained for 3 to 4 weeks at 24°C /22°C, 16/8 h day/night condition to allow silencing to occur.

### Extraction of Plant Total Proteins and Immuno-blotting

Each tomato and tobacco sample was homogenized in 300 μl 1× Laemmli buffer and then boiled for 5 min, followed by being resolved using 10% SDS-PAGE. Each Arabidopsis sample was homogenized in 300 μl protein extraction buffer (25 mM Tris-HCl, pH 7.5, 150 mM NaCl, 5% glycerol, 0.05%Nonidet P-40, 2.5 mM EDTA, 1 mM phenylmethylsulfonyl fluoride and 1× complete cocktail of protease inhibitors). The concentration of total proteins was determined using protein assay agent (Bio-Rad). Extraction of each sample containing 20 μg proteins was added with 2× SDS protein loading buffer and boiled for 5 min, then resolved by 10% SDS-PAGE. The immuno-blottings were performed with appropriate antibodies: anti-FLAG (Sigma), anti-HA (Sigma), anti-MYC (Santa Cruz), and anti-Ub (P4D1) (Santa Cruz).

### Bacterial Population Assay

The bacterial population assay was conducted as described previously (Katagiri et al., 2002; Nguyen et al., 2010). Briefly, for assaying the DC3000Δ*hopQ1-1* growth, *N. benthamiana* plants about four weeks after VIGS infection were first vacuum infiltrated with *P. fluorescens* 55 (*P. flu*55) by submersion of the aerial parts of the plant in a suspension of *P. flu55* (5 × 10^7^ CFU/mL) containing 0.002% Silwet L-77 and 10 mM MgCl_2_. The plants were then inoculated with *Pst DC3000ΔhopQ1-1* (2 × 10^5^ CFU/mL) in the presence of 0.002% Silwet L-77 and 10 mM MgCl_2_ by vacuum infiltration 7 h after the treatment with *P. flu55*. For assaying the growth of *Pst* strains DC3000 and DC3000*ΔhrcQ-U*, about four-week-old Arabidopsis or tomato plants about four weeks after VIGS infection were inoculated with the suspension of pathogen DC3000 (1 × 10^6^ CFU/mL) and DC3000*ΔhrcQ-U* (1 × 10^9^ CFU/mL) containing 0.002% Silwet L-77 and 10 mM MgCl2 by vacuum infiltration. In addition, the dipping method was also used for inoculation of tomato plants with DC3000*ΔhrcQ-U* (1 × 10^9^ CFU/mL). Inoculated plants were maintained in a growth chamber and monitored daily for symptom development. To assess bacterial populations, leaf discs were harvested from three to four plants of each treatment on day 3 and day 4 after the inoculation and ground, serially diluted, and plated to determine the amount of the bacteria grown as described (Zhou et al., 2017).

### Cell Death Suppression Assay

The cell death suppression assay was performed as previously described (Nguyen et al., 2010). The *P. fluorescens* 55 at the concentration of OD_600_ equal to 0.5 (~ 2.5 × 10^8^ CFU/mL), 0.1 (~ 5 × 10^7^ CFU/mL), and 0.015 (~ 7.5 × 10^6^ CFU/mL), respectively were used as the PTI inducer. The *Pst* strain DC3000 at the concentration of 2 × 10^6^ CFU/mL was used as the challenger in the assay. The challenge of PTI was conducted 7h after PTI induction. Appearance of cell death in the overlapping area, where both the inducer and challenger were infiltrated was assessed. Photographs were taken on the fourth day after infiltration of *Pst* DC3000.

### Accession Numbers

Sequence data of tomato, Arabidopsis, *N. tabacum, N. benthamiana*, wheat and soybean that were used in this article can be found in the GenBank data library based on the accession numbers: SlUBA1, Solyc06g007320.2.1; SlUBA2, Solyc09g018450.2.1; NbUBA1 Niben101Scf09017g00015.1; NbUBA2, Niben101Scf03202g13010.1; GmUBA3 (Glyma.02G229700), XP_003518319; GmUBA2 (Glyma.11G166100), XP_006591250; GmUBA1 (Glyma.14G196800), KRH17078; GmUBA4 (Glyma.18G058900), XP_006602078; TaUBE11, P20973.1; TaUBE12, P31251.1; TaUBE13, P31252.1; AtUBA1, AT2G30110; AtUBA2, AT5G06460; AtUBC32, AT3G17000; AtUBC33, AT5G50430; AtUBC34, AT1G17280; SlUBC32, KY246924; SlUBC33, KY246925; SlUBC34, KY246926; SlHRD1A, Solyc03g096930.2.1; SlHRD1B, Solyc06g072790.2; SlHRD3A, Solyc03g118670.2.

## Acknowledgments

We thank Christian Elowsky and You Zhou (Biotechnology Center, University of Nebraska-Lincoln) for helping with the confocal microscope and G.B. Martin for critical reading of the manuscript. We are grateful to Harkamal Walia for sharing the barley seeds for cloning of the MLO gene. We also thank the *Arabidopsis* Biological Resource Center for the *Arabidopsis* T-DNA insertion lines. This work was supported by start-up funds from the University of Nebraska-Lincoln, the U.S. Department of Agriculture National Institute of Food and Agriculture (grant no. 2012-67014-19449), and the National Science Foundation (grant no. IOS-1645659) to L.Z.

## Author Contribution

B.Z. designed experiments, performed most experiments, analyzed data, and wrote the manuscript. X. C. cloned the two tomato E1 genes, purified recombinant E1 and some E2 proteins and conducted the thioester assay to measure specificities of the tomato E1s in charging some E2 enzymes. C.W. examined the accumulation of MLO-12 in *N. benthamiana* UBC32i and UBC33/34i transgenic lines and the Arabidopsis *ubc32*, *ubc33*, and *ubc34* single, double and triple mutants, and performed the flg22 and elf18-triggered seedling growth inhibition assay. Y. Z. examined the subcellular localization of tomato SlHRD1A and SlHRD1B. L.Z. designed experiments, analyzed the data, and edited the manuscript.

